# Species Richness and Speciation Rates for all Terrestrial Animals Emerge from a Synthesis of Ecological Theories

**DOI:** 10.1101/2022.10.26.513952

**Authors:** Lucas D. Fernandes, Rogier E. Hintzen, Samuel E. D. Thompson, Tatsiana Barychka, Derek Tittensor, Mike Harfoot, Tim Newbold, James Rosindell

## Abstract

The total number of species on earth and the rate at which new species are created are fundamental questions for ecology, evolution and conservation. These questions have typically been approached separately, despite their obvious interconnection. In this manuscript we approach both questions in conjunction, for all terrestrial animals, which enables a more holistic integration and generates novel emergent predictions. To do this, we combine two previously unconnected bodies of theory: general ecosystem modelling and individual based ecological neutral theory. General ecosystem models provide us with estimated numbers of individual organisms, separated by functional group and body size. Neutral theory, applied within a guild of functionally similar individuals, connects species richness, speciation rate and number of individual organisms. In combination, for terrestrial endotherms where species numbers are known, they provide us with estimates for speciation rates as a function of body size and diet class. Extrapolating the same rates to guilds of ectotherms enables us to estimate the species richness of those groups, including species yet to be described. We find that speciation rates per species per million years decrease with increasing body size. Rates are also higher for carnivores compared to omnivores or herbivores of the same body size. Our estimate for the total number of terrestrial species of animals is in the range 1.03 − 2.92 million species, a value consistent with estimates from previous studies, despite having used a fundamentally new approach. Perhaps what is most remarkable about these results is that they have been obtained using only limited data from larger endotherms and their speciation rates, with the rest of the predictive process being based on mechanistic theory. This work illustrates the potential of a new approach to classic eco-evolutionary questions, while also adding weight to existing predictions. As we now face an era of dramatic biological change, new methods will be needed to mechanistically model global biodiversity at the species and individual organism level. This will be a huge challenge but the combination of general ecosystem models and neutral theory that we introduce here could be the only way to tractably achieve it.

## Introduction

Understanding speciation and how it varies between different groups is central to the interpretation of biodiversity patterns, and to our understanding of the fundamental processes that create biodiversity. Speciation, in turn, is conceptually linked to counts of the species it generates, providing an important gauge for the diversity of life. With planet Earth’s current trajectory towards a potential sixth mass extinction event (Barnosky et al., 2011; De Vos et al., 2015; Penn and Deutsch, 2022), the ability to estimate global species richness, including the many undescribed species, is all the more relevant to provide a baseline for what we have to lose.

A wide range of methods for estimating speciation rates exist. The majority of these focus on lineage-based models that are fitted to the fossil record and molecular phylogenetic data (Jetz et al., 2012; Pyron and Burbrink, 2013; Rabosky et al., 2014). Although new methodologies have been developed to take into account time, diversity, and trait dependences on speciation rates (FitzJohn, 2010; Pyron and Burbrink, 2013; Herrera-Alsina et al., 2019), these approaches are still limited by the availability of data on extinct and extant taxa. Moreover, speciation rates are inextricably connected to extinction rates and estimation of the latter is challenging, particularly in cases where there is no fossil data of now extinct species to inform methods (Rabosky, 2010). Recent work has also suggested that the inference models for speciation and extinction rates present intrinsic methodological limitations, leading to infinite indistinguishable diversification scenarios of speciation rates for the same time-calibrated phylogenies (Louca and Pennell, 2020).

One of the fundamental features of our planet, global species richness, including undescribed species, is a great challenge to estimate. Despite the potential for a conceptual link to the question of speciation rates, the question of global richness has been approached largely independently. One reason for the difficulty is that limited sampling has precluded the direct estimation of the total number of extant species. Indirect estimation of global richness has relied on the opinion of taxonomic experts (Gaston, 1991; Stork, 1993), inferences based on macroecological patterns (Raven, 1985; May, 1988; Hawksworth, 1991), and trends in taxonomic data accumulation (Mora et al., 2011).

However there is much controversy about the underlying assumptions of such approaches (Gaston and Blackburn, 2000; Bouchet and Duarte, 2006) and the large inherent uncertainty means the question of total diversity of species on earth remains open.

Given the methodological difficulties of estimating speciation rates and patterns of species richness using statistical inference, a promising alternative would be to obtain these estimates via mechanistic models of biological diversity. Indeed, there are many reasons to prefer mechanistic models over statistical inference or correlative models, especially where there is a lack of data (Grimm, 1999; Yates et al., 2000). By explicitly representing the underlying processes that affect the biodiversity units of interest, mechanistic models allow the exploration of novel ecological conditions and the generalisation of scenarios across data shortfalls (Connolly et al., 2017). Importantly, they also have emergent behaviour, making predictions that do not follow trivially from the original model assumptions. Such emergent predictions across multiple dimensions of ecology enable a stronger test of the model’s ability to capture underlying processes.

In this manuscript we combine the benefits of two very different mechanistic approaches: General Ecosystem Models (Harfoot et al., 2014) and Ecological Neutral Theory (Caswell, 1976; Hubbell, 2011). The Madingley Model is a General Ecosystem Model of global heterotrophic life (Harfoot et al., 2014), that makes predictions of, among other things, the numbers of individual organisms and demographic rates for different functional groups based on fundamental ecological assumptions. Neutral Theory (Hubbell, 2011), on the other hand, by ignoring functional differences and focusing primarily on species, provides a tractable approach to connect species richness, speciation rates and number of individual organisms within individual functional groups. The potential synthesis of these approaches could help to relax their individual assumptions, and build a more unified theoretical ecology structure that integrates both functional group and species-focused paradigms. The concept follows previous work based on the idea of nesting neutral models within a niche-based model of some kind (Chisholm and Pacala, 2010; Chisholm et al., 2016).

We implement a novel approach, building a mechanistic pipeline that connects the Madingley and neutral models. The Madingley model allows us to estimate numbers of individual organisms globally for functional groups of terrestrial endotherms and ectotherms. Neutral Theory allows us to connect number of individuals to speciation rates and species richness within any given functional group. By linking these models, we are able to estimate speciation rates for functional groups with known species richness.

Conversely, we can estimate species richness for functional groups with given speciation rates. We first apply the pipeline to well known groups of endotherms based on empirical species richnesses. We then extrapolate these patterns to estimate speciation rates for ectotherms. Finally we apply the pipeline in reverse to recover the expected species richness of ectotherms, including all yet to be described species.

Our aim with this work is to demonstrate a proof of concept for a novel synthesis of mechanistic ecological theories. Then to show their utility as a way to calculate speciation rates and estimate global richness for all terrestrial ecotherms and endotherms.

## Materials and methods

First, we ran the Madingley model and gathered data from it including total number of individual organisms in each functional group. Next, we developed individual-based ‘life history’ simulations of reproduction, maturation and death to acquire generation lengths of each functional group, based on the Madingley simulations. We gathered empirical data on global species richness of terrestrial endotherms and use it with analytical approximations of a neutral model, and the numbers of individual organisms from the Madingley model, to predict speciation rates per capita per generation for each functional group. We then convert these to rates per species per million years using the generation length data. Finally, we extrapolate these speciation rates for ectotherms and use the neutral model and numbers of individuals from the Madingley model again, but this time to recover species richness. 1.

### The Madingley Model

The Madingley model divides global-scale marine and terrestrial environments into two-dimensional grid cells. Each grid cell contains a mixture of individual organisms that may interact in complex ways and disperse to other grid cells (Harfoot et al., 2014).

Heterotrophic life is divided into nineteen functional groups that differ in their thermoregulation (endotherm or ectotherm), reproductive strategy (iteroparous or semelparous) and diet class (herbivore, omnivore or carnivore). Further ecological generalisations are used within each of these groups: a continuously varying body mass trait dictates the metabolic scaling, movement and trophic interactions of an organism across distinct juvenile and adult life stages. Ecological interactions are governed by fundamental mechanisms such as size-structured predation (Williams et al., 2010) and type-III Holling functional responses (Holling, 1959). Since tracking each individual in a global-scale implementation of the model would be computationally infeasible or arguably even impossible (Purves et al., 2013; Harfoot et al., 2014), identical individual organisms within a given local community are grouped together in ‘cohorts’. Each cohort is characterized by its categorical functional group and by its continuous body-size trait. All the individuals in the same cohort reproduce, mature, disperse or die as a group, with either the same process rate or probability of dying applying to all individuals within any given cohort. This kind of approach may be the only way to make a global individual-based mechanistic models tractable (Purves et al., 2013).

### Probing the Madingley model simulations for data

As it is computationally restrictive to collect data independently on every cohort in the model, we divided endothermic heterotrophic life in the Madingley model into discrete bins based on body mass. Starting from the minimum body mass, for each functional group, successive bin boundaries differ by a factor of two. Using herbivorous endotherms as an example, the first bin includes masses from 1.5 g to 3.0 g, the second from 3.0 g to 6.0 g, and so on. Thus, pairs of individuals whose body size differs by a factor of more than two cannot be in the same bin. This logarithmic binning gives a higher resolution on the small end of the body mass distribution, where most individuals lie, and is more natural for investigating allometric scaling. The same procedure was conducted separately for each of three diet classes of endotherms: herbivore, carnivore and omnivore; we do not refer to these as trophic levels because all three diet classes vary over orders of magnitude in size. Each size bin, within each functional group, is hereafter referred to as a ‘guild’. Whenever a mass value for a guild is used, it represents the midpoint body mass (in arithmetic space) of the bin in which that guild sits. We collected per capita rates of maturation, death and dispersal between adjacent cells, as well as the number of births and total abundance. These rates were collected separately for each guild, for each month of the annual cycle, and for each terrestrial cell of the Madingley model.

We ran the Madingley model at 2-degree resolution for a 100-year burn-in period, and then for a further 100 years for data collection. This length of time is sufficient for these models to be stable (Harfoot et al., 2014). We used default settings for the Madingley model, which did not factor in anthropogenic habitat loss or climate change (Harfoot et al., 2014). During the data collection period, the relevant demographic rates were recorded as running means. We performed fifteen data-collection runs, and used an ensemble mean across the 100-year data collection period and across all fifteen simulations in our downstream analyses.

### Individual-based simulations and generation time

It is necessary to know generation length in years for each guild in the Madingley simulations. This is because the neutral theory based speciation rates are a probability per birth event whilst we wish to compare this to rates per species per million years. In order to obtain an accurate estimate of the emergent generation times within the Madingley model, we performed a set of independent, individual-based simulations on a spatially explicit grid using process rates collected from the Madingley simulations. To do this, for each guild, in each grid cell, *I* = 1000 independent adult individuals were simulated from month to month. The starting month was chosen randomly but in proportion to the fraction of juveniles that mature into adults in each month. For example, if half of juveniles mature into adults in May and the other half in June then half of the individual based simulations would likewise start in May and the other half in June. In a grid cell, any given ‘focal’ adult either progressed to the following month, or died based on the mean mortality rate measured from the Madingley model simulations for that guild, location and month. Adult lifespan, in months, was recorded for any individuals that die. At each time step, the surviving focal adult may disperse to an adjacent grid cell depending on a random draw against the recorded mean dispersal rate out of that grid cell. The focal adult also stochastically produces offspring based on a Poisson process in each month. The number of offspring was given by the average number of births in the Madingley model, taking into consideration spatial and temporal heterogeneity in rates. Each offspring was then also simulated, in a similar manner to the adults, from the month and location of birth, until reaching reproductive maturity. Offspring were subject to juvenile mortality and dispersal checks analogous to those of adults, but based on juvenile-specific rates collected separately from the Madingley model simulations. Juveniles are not reproductively active and instead they had a stochastic opportunity to mature based on the maturation rate for the grid cell they occupy in each specific month of the year. The number of time steps to maturity, in months, was recorded if the individual offspring did not die before maturing. We calculated generation length for each guild in month-long time steps that we then scaled to years. The generation length is the difference between the date of birth of an individual and the average date of birth of its offspring. Reproduction events happen throughout adult life so the average time of offspring birth is the midpoint of the part of an individual’s life when it is a reproductive adult. Generation length can thus be calculated as the sum of two components: the mean time between birth and maturity of a juvenile, plus half of the time between maturation and death of an adult.

### Metacommunity richness with protracted speciation

After an initial divergence between two populations of a given species, it might take several generations before they can be considered separate species. One way to include this mechanism in a neutral model is to consider an additional parameter *τ*, giving the duration of speciation as a number of generations. This mechanism is known as protracted speciation (Rosindell et al., 2010). The per capita speciation rate, *ν*, may thus be smaller than the speciation-initiation rate, *μ*, and the two are related by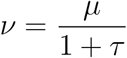 (Rosindell et al., 2010).

We obtain an analytical solution (see Supplementary Material) for the metacommunity richness of a non-spatial neutral model with protracted speciation given by:

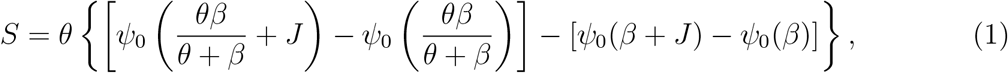

where *J* is the community size, 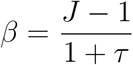, and 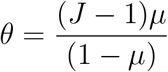. Note how *θ* depends on the speciation-initiation *μ*, instead of the per capita speciation *ν*.

Whilst this solution is for a non-spatial model - we show by simulations (see supplementary material) that it also works as an approximation for the global species richness in a spatially-explicit model with demographic rates parameterised from the Madingley model.

### Endotherm Empirical Data

We obtained empirical data for species richness of terrestrial endotherms in each guild using the EltonTraits 1.0 database of dietary and habitat requirements for birds and mammals (Wilman et al., 2014). We filtered the available data to ensure it was comparable to the ecology of simulated organisms in the Madingley model. Specifically, we removed all mammals that had “marine” foraging stratum category, and all of those whose diet consisted of more than 10% fish, as this resource is not explicitly modelled in the terrestrial-only simulations. To account for species that use freshwater wetlands and waterways, not explicitly modelled in Madingley, we also excluded species with any reference to “aquatic”, “freshwater”, or “glean” on the comments for foraging stratum. For birds, we excluded those assigned “Pelagic specialist”, those that had “Vertebrates and Fish and Carrion” as dominant diet category and those whose diet consisted of more than 10% fish. We also removed birds whose prevalence of foraging on water surfaces (summing “below surface” and “just below surface”) was greater than 50%. From a database with a total set of 9995 birds, we were left with 8677 (87%), and from 5403 mammals, 5084 (94%) remained.

We then categorised the remaining comprehensive list of terrestrial endotherms into three dietary categories: herbivore, carnivore, and omnivore. A species was considered a herbivore if the sum of the diet prevalence of fruit, nectar, seed or other plant resources was equal to 100%, a carnivore if the sum was equal to 0%, and an omnivore for everything in between.

### Speciation rates per species per million years

With the empirical values for endotherm richnesses and estimates of endotherm abundances from the Madingley model, we estimated speciation rates for each diet class at each interval of body mass. For each guild, we numerically solved equation 1 for the speciation-initiation rates *μ*, with *S* for empirical richness and *J* for Madingley abundance, and *τ* as a free parameter.

The speciation-initiation rates correspond to probabilities of initiating speciation per individual per birth/death event, thus not directly comparable to speciation rates estimated using phylogenetic methods, usually presented in units of per species (or per lineage) per million years. To obtain rates with comparable units, we first calculate speciation-completion rates 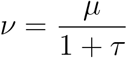, then perform the following transformation:

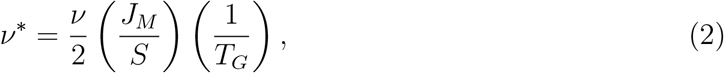

where *ν* is the speciation-completion rate (per individual per birth/death event), 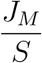 is the average number of individuals per species (considering simulated individual abundances and empirical species richnesses), and *T*_*G*_ is the generation time (in million years), obtained from the individual-based simulations using demographic rates probed from the Madingley simulations. The factor of 2 arises because we are modelling overlapping generations. As a result, the generation length is half the adult lifespan; relatedly, half of living adults are post reproduction and belong to an older generation (Rosindell et al., 2010). Equation 2 guarantees that *ν*^∗^ expresses the speciation rate for a given guild in unit of per species per million years.

### Ectotherm richness

The method that we applied to estimate speciation rates for endotherms, as functions of body mass and diet class, can be reversed to infer ectotherm richnesses. First, we fitted power-law functions separately for each diet class, to per species per million years speciation rates, and to generation times, as functions of body mass. We then extrapolated the results to obtain estimates of these quantities for smaller values of body mass, in particular those corresponding to the smaller body size guilds within ectotherm functional groups. For each guild of ectotherms, we have Madingley simulated abundances, the extrapolated values of speciation rate *ν*^*^, and generation time *T*_*G*_ from our individual-based simulations. We thus obtain, for each guild, the speciation rates per individual per birth/death event, *ν*, as a function of the unknown species richness, *S*, by inverting equation 2. Thus, for each body mass guild, we obtain an expression for *ν* in terms of *S* that can be substituted into equation 1, allowing us to numerically solve the result for *S*. The resulting *S* provides an estimate of total species richness for that ectotherm guild.

## Results

### Madingley abundances and empirical richnesses

The total abundance of herbivores and omnivores in the Madingley simulations shows a decreasing trend with increasing body mass, for body masses above 10^3^ *g* (left and right panels, first row - Fig. 2). Herbivores further show a slight interior peak in abundance at just under 10^6^ *g* despite an overall decreasing trend.

**Fig. 1.**
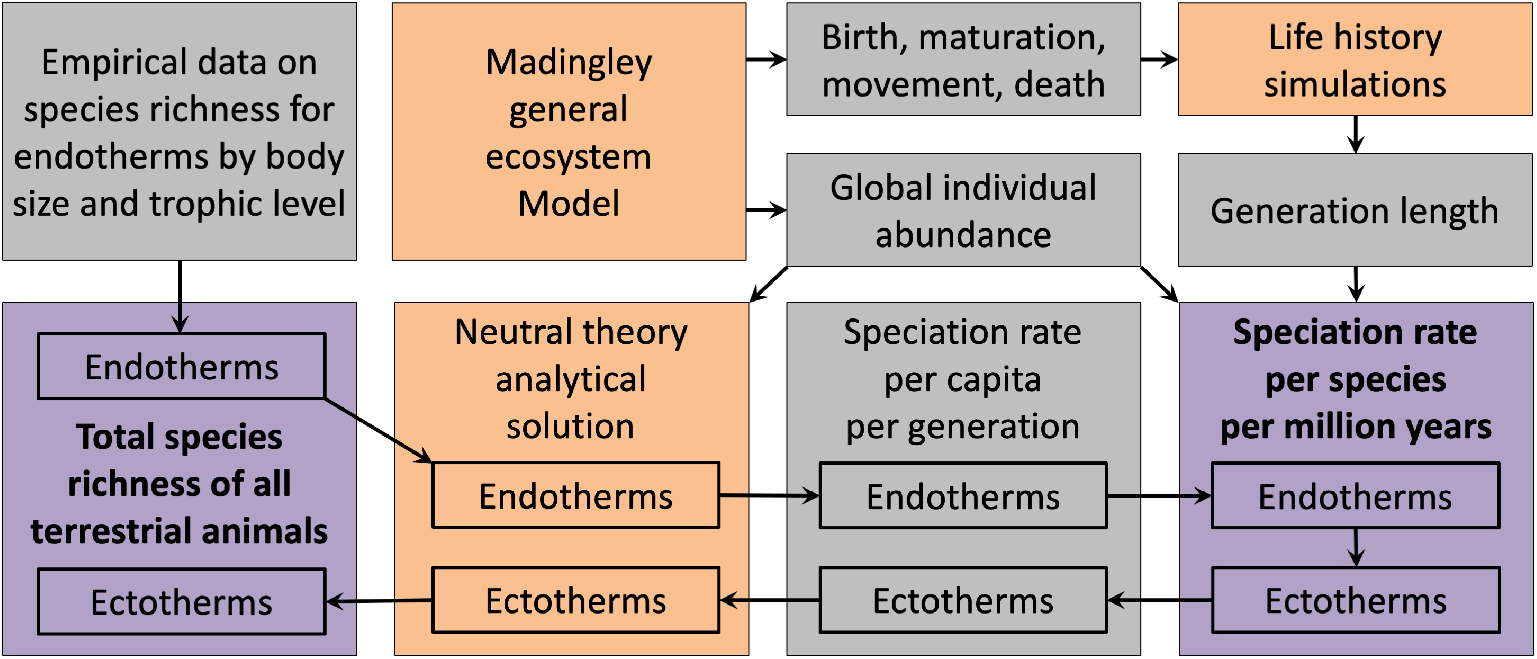
Conceptual overview of the methods and analysis pipeline. Purple boxes correspond to our main results, orange boxes show simulations and analytical models, grey boxes show other forms of input and intermediate data products that are used in downstream analysis. The bottom four boxes represent a pipeline that can convert from species richness and number of individuals to speciation rate per species per million years. The same pipeline can be run in reverse in order to recover species richness if the speciation rate and number of individuals are known. The full analysis shown in this diagram is repeated for a full range of functional groups (e.g. herbivore, carnivore and omnivore functional groups in terrestrial endotherms) as well as across a wide range of body size classes within each functional group.

**Fig. 2.**
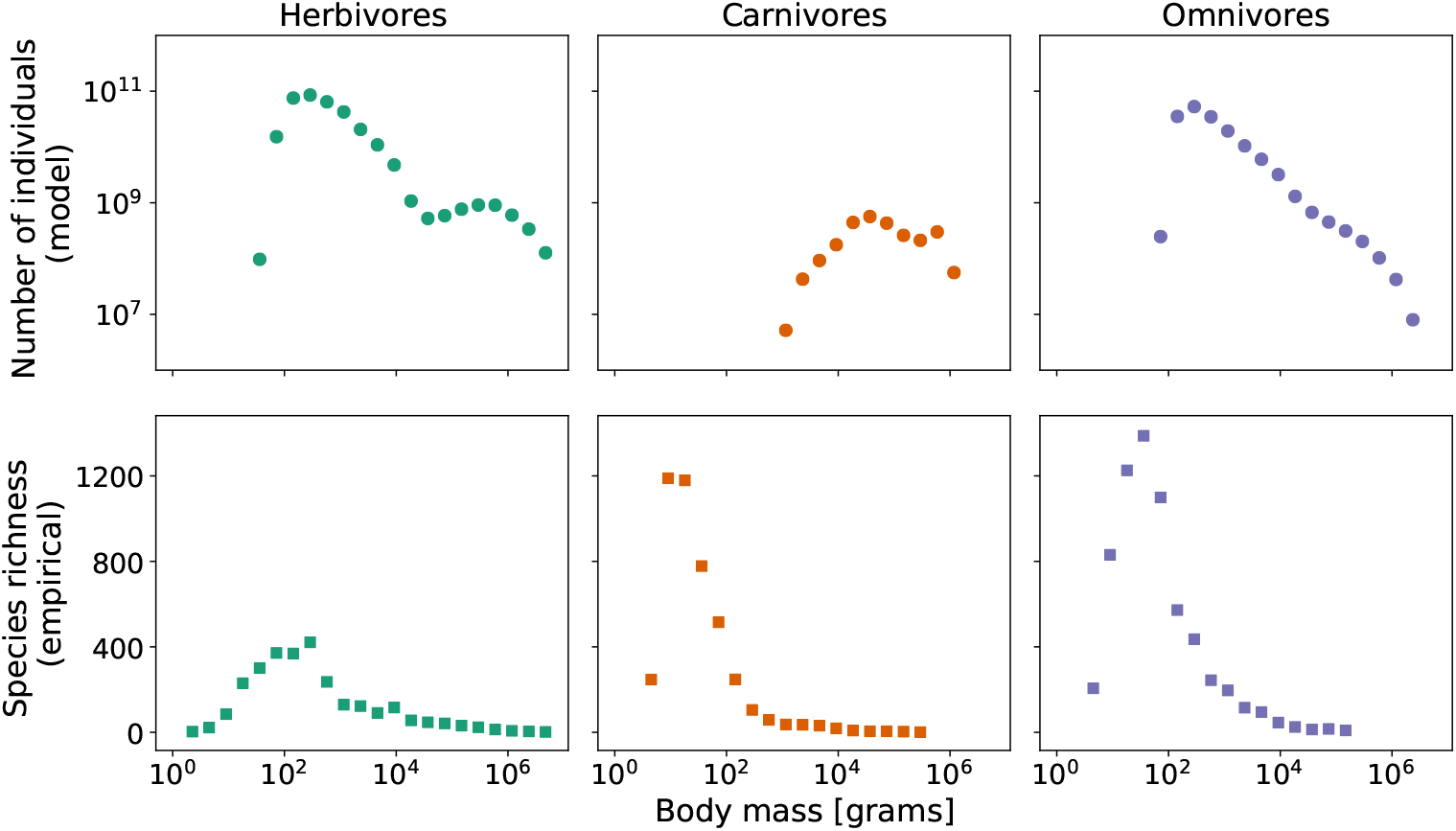
Average global abundances (in number of individuals) from the Madingley model at equilibrium for terrestrial endotherms (first row). Total empirical species richness (second row) for terrestrial endotherms. All shown as a function of body mass: the midpoint of the body mass interval corresponding to each guild. Separate results are shown for three diet classes: herbivores (green), carnivores (orange), and omnivores (purple).

Terrestrial endothermic herbivores and omnivores with body masses below 10^2^ *g* were scarce or absent in the Madingley simulated data. Similarly, terrestrial endothermic carnivores with body masses below 10^3^ *g* were absent in our simulations (central panel, first row - Fig. 2). Abundances of carnivores peak at a body mass of 10^5^ *g*, with a slight tendency of decreasing abundances at smaller and larger body masses.

Empirical species richness of endotherms, based on the EltonTraits database (Wilman et al., 2014) after our filtering process, peaked at a body mass of approximately 10^2^ *g* for herbivores and omnivores, and at a body mass of around 10 *g* for carnivores (second row - Fig. 2). Numbers of species dropped sharply either side of the peak. The Madingley simulations supported carnivores and omnivores with large body masses (above 3 · 10^5^ *g*) that are absent in the empirical species data.

### Protracted speciation rates and generation times

For all diet classes, patterns of speciation-initiation rates as a function of body size were largely independent of the values of *τ* (Fig. 3). Speciation-initiation rates, for all three diet classes, show a sharp decline with increasing body size for organisms of extremely small size. After this, the herbivore speciation rate increases with body mass to form an interior mode between 10−3 *g* and 10−4 *g*, whilst the omnivore speciation rate increases subtly with increasing body size, and the carnivore speciation rate remained flat.

**Fig. 3.**
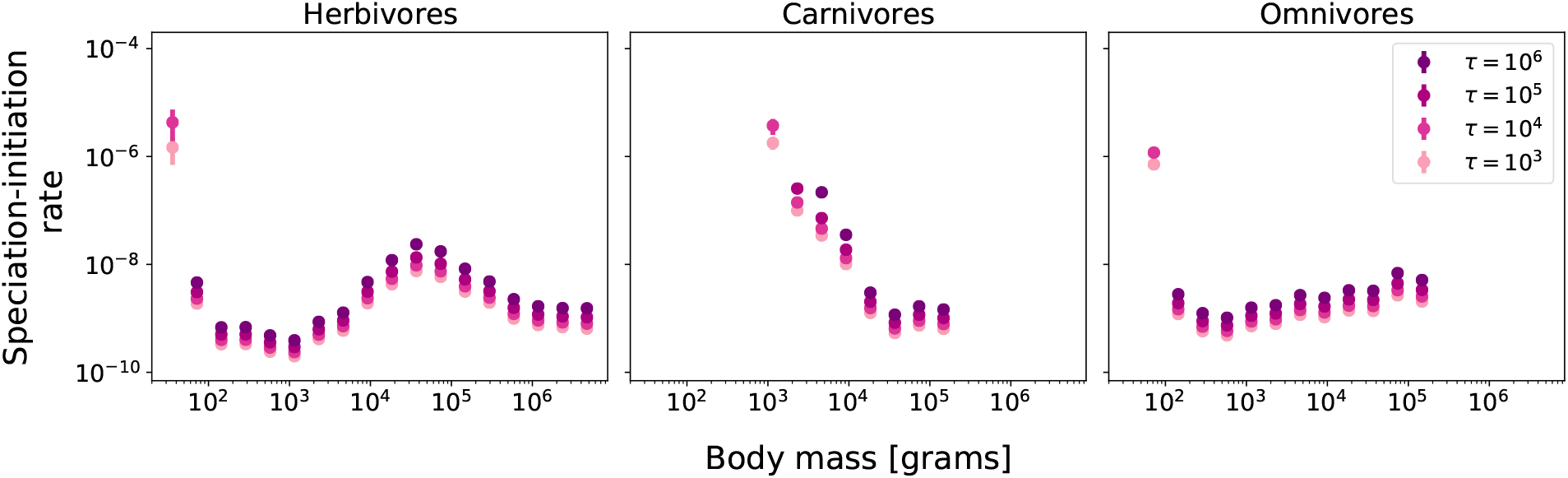
Numerical estimates of speciation-initiation rates (protracted speciation neutral model) for herbivore (left), carnivore (center), and omnivore (right) terrestrial endotherms, as a function of body mass. Results are shown for four different durations of protracted speciation: *τ* = 10^3^, 10^4^, 10^5^, and 10^6^. Results are obtained as average values of speciation-initiation over 15 different samples of Madingley abundances and error bars correspond to one standard error of the mean. Often the error is smaller than the size of the point symbol and in these cases it is not visible.

Increasing values of *τ* led to increasing estimates of speciation-initiation rates, for all guilds in each of the dietary groups. For each diet class, some of the values of speciation-initiation rates for the smallest body mass guilds (*τ* = 10^5^ and *τ* = 10^6^) are not shown because equation 1 presented no numerical solution for *μ* (see Supplementary Material).

Estimates of speciation rates were limited for small-bodied groups (particularly below 100 *g* for all groups) due to the absence of terrestrial endotherms within those body mass intervals in the Madingley model simulations (Fig. 2). For carnivores, only guilds with body mass above 10^3^ *g* show stable results for equilibrium abundances. Since the peaks for empirical richnesses for carnivores and omnivores are located between 10 and 100 *g*, speciation rates for a significant portion of the global biodiversity for these groups could not be evaluated by this method.

Estimated values of the generation times for all the guilds of terrestrial endotherms are below 20 months (Fig. 4). Across all guilds for which the three diet classes are present, omnivores show the highest generation times, followed by herbivores, and lastly carnivores. Carnivores and large-sized herbivores (above 10^4^ *g*) are reasonably consistent with the theoretical allometric scaling for generation times, which predicts an increase of generation length with body mass as a power-law with exponent 0.25 (Savage et al., 2004). For omnivores, the comparatively reduced slope indicates slower increase of generation time with body mass.

**Fig. 4.**
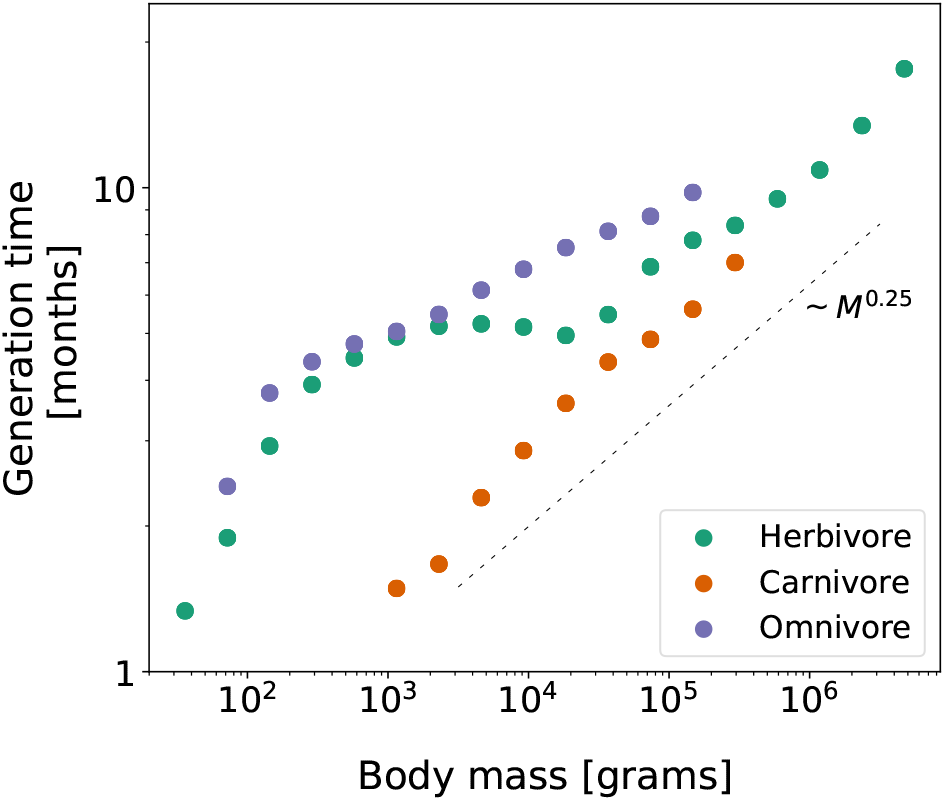
Generation times for terrestrial endotherm guilds. The times were estimated from individual-based simulations using stationary demographic rates in the Madingley model, as a function of body size. Results are shown for herbivores (green), carnivores (orange), and omnivores (purple). The dashed line shows a reference slope for the allometric scaling ∼ *M* ^0.25^ with arbitrary intercept.

### Speciation rates for terrestrial endotherms

Speciation rates, converted to units of per species per million years, show a consistent pattern of decline with body mass in the three diet classes of terrestrial endotherms (Fig. 5). In the case of the herbivore diet class, there is additionally an almost flat relationship between body size and speciation rate at intermediate body sizes. For most guilds speciation-initiation varies in a range smaller than one order of magnitude for all the assessed values of *τ* (see Fig. 3). As a result, an inversely proportional relation approximately holds between *τ* and speciation rate per species per million years.

**Fig. 5.**
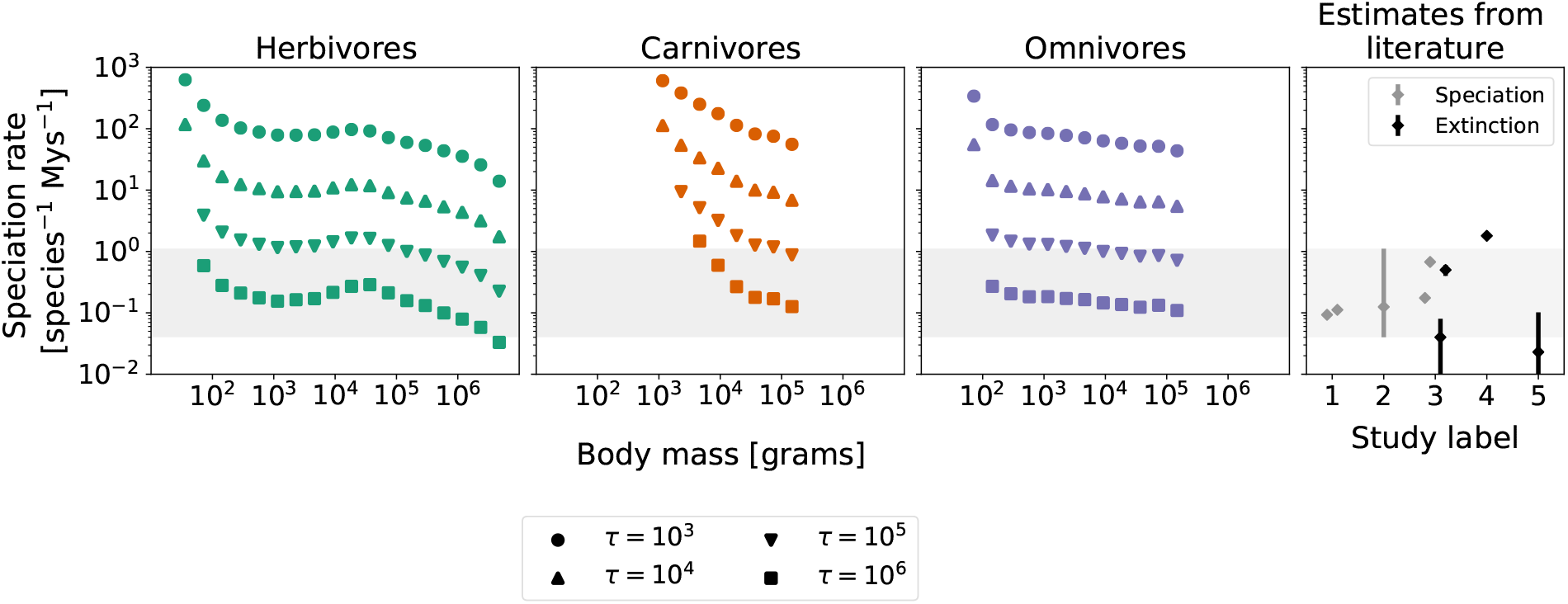
(First three panels) Speciation rates per species per million years as a function of body mass for three diet classes of terrestrial endotherms. For each class, the rates are calculated for four distinct values of speciation-completion time: *τ* = 10^3^, 10^4^, 10^5^, and 10^6^. Rates for each guild correspond to mean values over 15 estimates, using stationary abundances from different Madingley simulations. Values for the standard errors of the mean are very small compared to the mean values, therefore error bars are not visible. (Rightmost panel) Estimates of speciation and extinction rates (gray and black diamonds, respectively, in units of event per species per million years) obtained from previous studies, labeled as follows: (1) Rolland et al. (2014), (2) Rabosky et al. (2015), (3) Weir and Schluter (2007), (4) Barnosky et al. (2011), and (5) De Vos et al. (2015).

The comparison between the three diet classes also shows that, while herbivores and omnivores present similar values of speciation rates for the same body size group, speciation rates for carnivores are often higher, for the interval where the rates could be estimated for the three groups (approximately between 10^3^ *g* and 10^5^ *g*). The difference was particularly clear for smaller-bodied carnivores (see Supplementary Material).

Direct comparison is also possible with estimates of speciation rates for birds and mammals from previous works (see Fig. 5 (gray diamonds, rightmost panel). The speciation rates we obtain with our method are consistent with the range of values obtained from the literature through entirely different methods, particularly for values of *τ* equal to 10^5^ and 10^6^.

By using equation 1 to estimate speciation rates, we implicitly assume that the empirical patterns for richness correspond to equilibrium richness of the neutral model with protracted speciation. In a neutral model, net zero diversification rate is reached at equilibrium, when speciation rates equal extinction rates. Thus, the values we obtain for speciation rates as functions of body mass in Fig. 5 also correspond to extinction rates (under the assumptions of our model). Direct comparison with estimates of baseline extinction rates from the literature (black diamonds, rightmost panel - Fig. 5) also reveals that extinction rates obtained via our method are consistent with those obtained from phylogenetic analysis, for values of *τ* = 10^5^ and 10^6^.

### Mechanistic estimates of ectotherm richness

Fitted plots and parameter tables for the power law functions of speciation rate per species per million years can be seen in the Supplementary Material. We have estimated richnesses for ectotherms, as functions of body mass and diet, by extrapolating these functions for *τ* = 10^5^ and 10^6^, the two values of speciation-completion time which are more consistent with previous estimates from phylogenetic approaches (see Fig. 5). The results for ectotherm richnesses can be seen in Fig. 6. As a general trend, species richness decreases with increasing body mass for all groups, at least for larger values of body mass. At the small body size end of the spectrum there is a relatively brief band where this relationship is reversed. For semelparous ectotherms, while herbivores and omnivores present similar values of abundance at each body mass bin, values of speciation rates for herbivores are greater than the ones for omnivores (for the same values of *τ*) in the body mass range from 10^−2^ to 10^4^ *g*. For semelparous carnivores, although values of abundance are orders of magnitude lower in comparison to herbivores and omnivores, speciation rates are higher, for values of body mass below 10^5^ *g*. Summing the richnesses of all body mass intervals for the three diet classes, we have total global richnesses for semelparous ectotherms ranging from 9.97·10^5^ to 2.75·10^6^ species, and for iteroparous ectotherms ranging from 1.75·10^4^ to 1.56·10^5^ species (the smallest values corresponding to *τ* = 10^6^ and the highest, to *τ* = 10^5^). The total number of terrestrial animals, including our predicted numbers of ecotherms, and our empirically derived number of 13, 703 endotherms, was between 1.03 *−* 2.92 million species, depending on the value of *τ*.

**Fig. 6.**
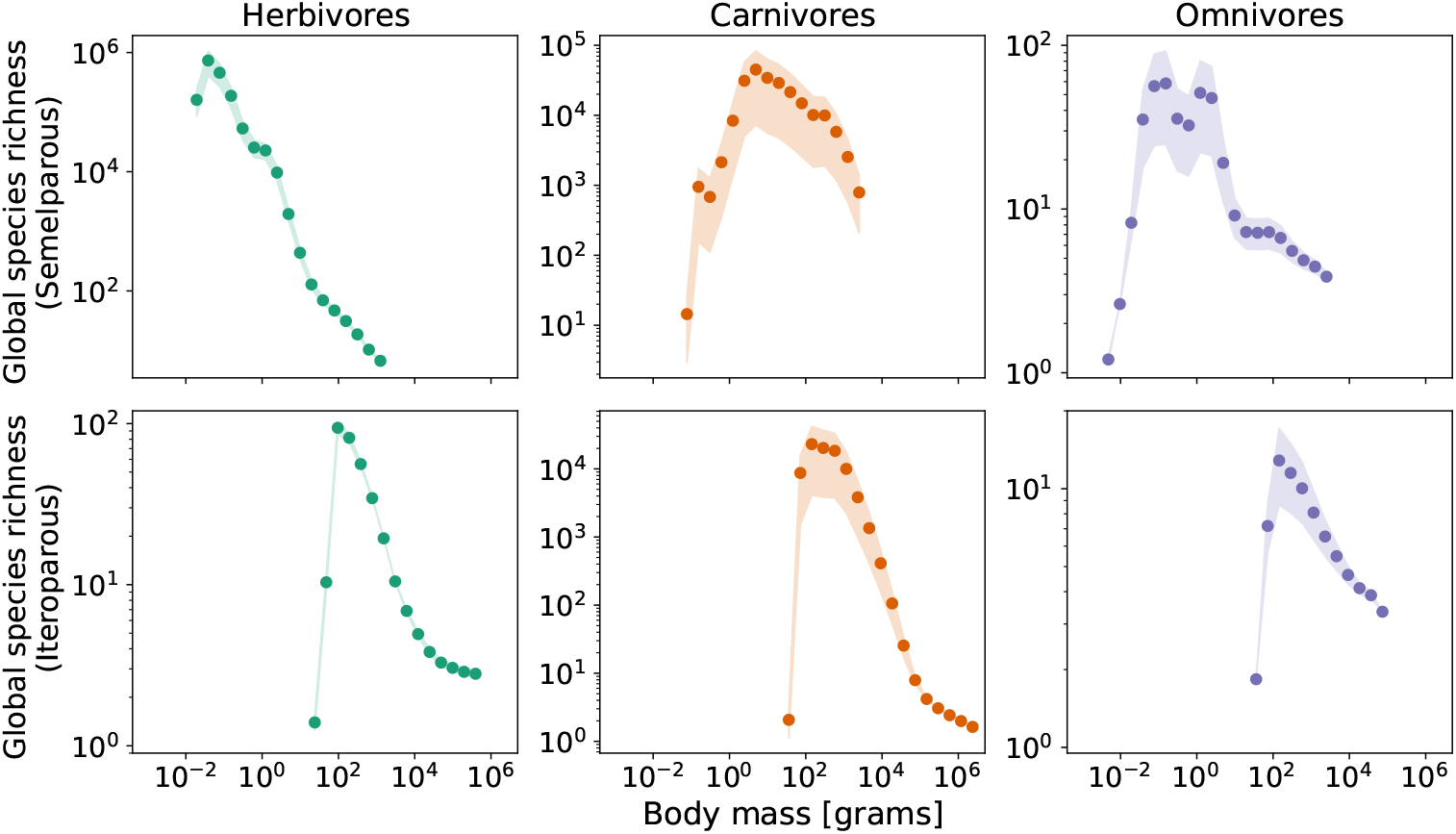
Global species richnesses for terrestrial ectotherms, as functions of body mass and diet. Results are shown for two reproduction strategies, semelparous (first row) and iteroparous (second row), following the definitions of functional groups in the Madingley model. Shaded regions correspond to the range of estimated richnesses for each body mass for the two values of speciation-completion times used (*τ* = 10^5^ and 10^6^), while circles correspond to the mean values for these ranges.

## Discussion

In this study, we sought to connect two previously unconnected areas of theory to integratively address two questions that have so far been approached independently. The ‘Madingley model’ (Harfoot et al., 2014) is a first-of-its-kind general ecosystem model, able to make mechanistic global-scale predictions about broad functional groups of organisms on land and in the ocean. One key limitation to the Madingley Model, however, is that it omits species identities and so cannot make predictions about species richness or speciation. In contrast, ecological neutral models (Hubbell, 2011) don’t incorporate the functional differences that are captured by the Madingley Model, but focus instead on making predictions about species, including speciation and extinction rates. By combining the two, we introduce a body of mechanistic theory that operates tractably at the global level and also includes the species concept. In this manuscript, we applied our approach to estimate speciation rates of terrestrial endotherms as a function of diet class and body size, and to estimate the total diversity of terrestrial ectotherms, including all undescribed species.

### Per-capita speciation and generation length

Our results for per-capita per-birth speciation-initiation rates *μ* represent an interim result from our analysis pipeline, relevant only in the context of neutral theory. It is still worthwhile to consider the mechanistic reasons behind the patterns in *μ* that we observe. These seem to be mostly arising from fluctuations in the numbers of individual organisms for different guilds predicted by the Madingley model. The hump in *μ* for intermediate body sizes of herbivores appears to correspond to slight dip in the total number of individual organisms at the same body size, as estimated by the Madingley Model. Similarly, our result that small carnivores diversify faster than herbivores or omnivores of the same body size coincides with there being fewer individual organisms without there being proportionately fewer species. Whilst the values of *μ* are not typically studied, it seems plausible to us that the empirical relationship with body size could be hump-shaped where there is a dip in numbers of individuals without a corresponding dip in numbers of species. This seems to be the case in our model.

To translate these per-birth speciation rates into a rate per species per million years required us to know both number of individuals per species and number of generations per million years. To derive the latter, we use generation lengths in years predicted by the combined use of the Madingley model and our individual-based simulations. Our estimated values of the generation times for all the guilds of terrestrial endotherms are below 20 months (Fig. 4). This is rather short, especially for guilds of large body size, and is consistent with the accelerated pace of life predicted for organisms in the Madingley model, as compared to empirical estimates (Harfoot et al., 2014). The relationship between generation length and body size did, however, show comparable results to a power law with exponent 0.25, consistent with previous expectations (Savage et al., 2004).

### speciation rate variability with body size and diet class

Our estimations of speciation rate per species per million years, as a function of body mass and diet class depend on the speciation-completion time, *τ* (the time it takes to form two separate species from two lineages that started to diverge). As a result, we have a range of speciation rates for the same diet class and body mass interval as *τ* varies (Fig. 5). However, while the resulting curves in figure 5 move up and down with *τ*, they do not change shape. Thus, the qualitative way that speciation rates vary with body size and diet class is independent of the value of *τ*, and thus remains a robust finding of our models.

In general, previous work across different taxa has reported mixed relationships that focus on diversification rate, species richness and body size (Orme et al., 2002; Wollenberg et al., 2011; Feldman et al., 2016). Overall, previous studies suggest that diversification may decrease with increasing body size, but may also be humped or with no relationship. Previous work has also approached the question of how diversification may vary with diet class, suggesting that herbivores diversify fastest, followed by carnivores and then omnivores (Price et al., 2012). What is noteworthy about previous work is the focus on net diversification, rather than on speciation, which is notoriously difficult to pick apart from extinction (Louca and Pennell, 2020). There is a general agreement that the answers remain unresolved. As a result, these previous works are neither supportive nor in contradiction with our results about speciation rate and its relationship with body size and diet. We did not find a single simple relationship between body size, diet and speciation rate, but certain results did stand out. For example, speciation rates were generally higher with smaller body sizes, a result that may be partly explained by differences in generation time (Etienne et al., 2012). We also found that smaller carnivores have higher speciation rates than herbivores and omnivores of the same body size. This may be because the higher values of *μ* for carnivores are not entirely cancelled out when we translate *μ* back into speciation rates per species per million years.

### quantitative speciation rates and background extinction rates

Overall, our speciation rates per species per million years, regardless of diet class and body size, are consistent with independent estimates for endotherms obtained from studies using different phylogenetic methods that do attempt to separate speciation from net diversification. For example, Rolland et al. (2014) obtain 95% credibility intervals for posterior distributions on speciation rates that range, approximately, from 0.085 to 0.101 for temperate mammals and from 0.107 to 0.116 for tropical mammals (study 1 – Fig. 5). Rabosky et al. (2015), working with bayesian inference on New World Birds phylogenetic data obtain per-taxon rates that range from 0.04 to 0.11 new species per million years (study 2 – Fig. 5). Weir & Schluter (Weir and Schluter, 2007) estimate intervals of speciation rates for birds and mammals varying with latitude, with estimated per-lineage per-million years rates of 0.15-0.2 at the Equator and 0.6-0.75 at 50^◦^ latitude, N or S (study 3 – Fig. 5). Overall, speciation rates for endotherms obtained from previous studies are within a range from 0.04 to 0.11, which corresponds to the range seen for *τ* between 10^5^ and 10^6^ (gray rectangles in Fig. 5). These values of *τ* are broadly consistent with expectations for the speed of segregation of lineages. This speed should certainly be less than the expected waiting time between successful speciation events, but should also not be small, which would correspond to a generally unrealistic scenario (Rosindell et al., 2010).

Under our neutral models, a dynamic equilibrium between speciation and extinction is assumed for any particular guild. This enables us to equate our speciation rates directly to extinction rates per species per million years and to a net diversification rate of zero.

Retaining the same range of *τ* between 10^5^ and 10^6^ the values we obtain for background extinction rates also present a reasonable correspondence with the range of extinction ratesobtained from previous studies, ranging from 0.0 to 1.8 extinctions per species per million years (or equivalently per million species per year, the more usual unit for background extinction rate) (Weir and Schluter, 2007; Barnosky et al., 2011; De Vos et al., 2015) (studies 3-5 – Fig. 5). Future work may consider relaxing the dynamic equilibrium assumption. In the context of this study, however, forcing net zero diversification rates provides a new perspective on speciation and extinction as well as how they vary with body size. Given the general difficulty in finding speciation rates, and background extinction rates, we suggest that both inferential methods and mechanistic models of different kinds are needed, and should ideally get consistent results, in order to provide convincing answers through a ‘mixed approach’.

### total species richness of terrestrial animals

The possibility of reversing the neutral model component of our pipeline to estimate species richness for ectotherms provides a novel approach to investigate biodiversity patterns of undersampled groups. Our results for estimated total species richness are sensitive to changes in speciation rates (per species per Mys) and, to a lesser extent, to values of ectotherm abundances (see Supplementary Material). This explains the large difference in predicted numbers of species between omnivores and the other two diet classes. Analogous reasoning can be used to interpret the differences between diet groups for iteroparous ectotherms. The positive correlation between richness and body mass for smaller values of body mass is not necessarily realistic, but is associated with the low abundances for small-bodied organisms in Madingley’s stationary distributions, a known limitation of the Madingley model (Harfoot et al., 2014), rather than of the way it is used in this application.

Overall, our estimated number of terrestrial species is between 1.03 and 2.92 million, dominated by guilds of ectotherms that likely correspond to arthropods. These results are consistent with current estimates of extant terrestrial arthropods of 7· 10^6^ species (Stork et al., 2018). Our estimated numbers of iteroparous ectotherms, ranging from 1.75· 10^4^ to 1.56· 10^5^ also compare well to extant reptile and amphibian fauna of 11570 and 8482 species, respectively (Uetz et al., 1995; AmphibiaWeb, 2022). This comparison, however, should be taken with caution because many iteroparous ectotherms will also be arthropods, and conversely many amphibians will not be terrestrial.

### limitations and potential future work

The version of the Madingley model from which we derived our data assumes a pristine world in which there has been no anthropogenic change. Given our aims are to understand historic speciation rates, background extinction rate (without anthropogenic species loss), this assumption seems justified. We calibrated our speciation rates to match presently extant terrestrial endotherms rather than the numbers of terrestrial endoterms that existed prior to anthropogenic influences. This will mean that our total numbers of species take into account anthropogenic change to some extent. The difference caused by this is probably relatively small for most body sizes since only a very small percentage of described species have so far been declared extinct and our pipeline only requires species numbers. Increasing numbers of endangered yet extant species would not affect our results. An exception would be large bodied species where hunting has caused a large proportion of extinctions due to anthropogenic change. This is likely why the Madingley model supported the existence of large bodied endotherms not matched by our empirical data 2. It would be possible to account for recently extinct species but this would require a comprehensive database of all recently extinct endothermic life, together with body size and diet data. Future work could more explicitly incorporate anthropogenic change and study its effects. For example, by changing the environmental and landscape input data in Madingley, according to projected scenarios of climate change. Similarly, habitat loss could be simulated via removal of a proportion of primary productivity from the landscape for use in food production and support of domesticated animals (Newbold et al., 2020).

Another limitation in our approach is the lack of small-bodied organisms in the stationary distributions in the Madingley model. This affects not only the estimation of speciation rates for endotherm body masses lower than 100 *g*, but also results in larger errors in the estimation of ectotherm richness by extrapolation of the endotherm rates. Stationary distributions of ectotherms in Madingley are also underrepresented in a large interval in comparison to the lower bounds allowed in Madingley for these groups (lower bounds for body masses of semelparous and iteroparous ectotherms in Madingley are 0.0004 *g* and 1 *g*, respectively). There are numerous possible drivers of the lack of abundance of small organisms in Madingley. For example, the lack of temperature-dependence for foraging parameters (Rall et al., 2012) may wrongly disadvantage small bodied organisms. The high relative metabolic costs may impose a disproportionately large additional energetic burden on small organisms (Sibly et al., 2012). Vegetation in Madingley is not vertically structured, so there is no distinct rainforest canopy for example, often it is vertical structuring that creates new niches favouring small bodied taxa (Krause et al., 2022). In the natural world, smaller organisms are often able to exploit flushes of high energy resources such as nectar, fruits and seeds, all of which are not explicit in the Madingley model. Another possibility is that since modelled endotherms in Madingley are active for one hundred per cent of a model time step and cannot seek metabolic refuge to limit their energy expenditure. They cannot optimise their energy budgets as natural organisms such as hummingbirds (Payne, 2010) and bats do by way of torpor and crepuscular behaviours. The effect of all these drivers on Madingley biodiversity distributions should be explored in future work.

Another promising line of investigation resides on the improvement of the performance of spatial neutral models. Here we used the analytical solution of non-spatial neutral models as an approximation to the global richness, with which we estimated speciation rates for endotherms and richnesses for ectotherms. Although it can be shown that the non-spatial analytical richness seems a reasonable first approximation for the global richness in a spatial model (see Supplementary Material), we know that abundance distributions in Madingley are strongly affected by spatial heterogeneities in availability of resources and species dispersal kernels. The main reason to use this approximation is the heavy computational cost of performing spatial neutral simulations (Thompson et al., 2020), with values of speciation rates and abundance needed to match the empirical patterns. In these spatial models, the introduction of optimisations such as Gillespie algorithms may lead to a significant gains in performance, allowing estimates of spatial richness distributions instead of only total global richness. Developments on this front could benefit not only our method’s estimates for speciation rates and undescribed species richnesses, but the field of ecosystem modelling as a whole, with mechanistic models playing a central role in producing theoretical and applied advancements.

## Conclusion

In this work we have synthesised two mechanistic models to connect previously disparate fields of ecological theory. The result is a new pipeline with synergistic models capable of making predictions about large-scale and partially observed phenomena, illustrated here with a study of species richness and speciation rates. Our method opens the possibility of exploring drivers of speciation and background extinction from a mechanistic approach; this opens up the possibility of making emergent predictions around other evolutionary processes and questions. The deeper potential of our approach is that it paves the way to incorporate anthropogenic change, and thus mechanistically predict anthropogenic extinctions with potential conservation applications. This line of investigation is hard to perform with traditional models based on phylogenetic inference, since they are not underpinned by a general ecosystem model or process-based ecological theory.

Perhaps what is most remarkable about our results is that both species richness and speciation rate estimates for all terrestrial animals have been obtained using only limited data from larger endotherms and their speciation rates, with the rest of the predictive process being grounded in mechanistic models. Our hope is that this work will pave the way for future general ecosystem model and neutral model combinations. Not just to apply this to the marine and freshwater realms, but also as part of an ongoing interest in general ecosystem models, that can now include predictions about species diversity. Whilst this does involve making many assumptions, it is likely the only approach that will lead to tractable process-based models of global species diversity from first principles. The addition of neutral theory to general ecosystem models introduces the species concept and brings the original vision of general ecosystem models one step closer to reality.

## Acknowledgements

We thank Andy Purvis and Drew Purves for very useful conversations and support. We thank the Leverhulme Trust for funding this research (grant number RPG-2018-069). Through JR and LDF this work is an output of the Georgina Mace Centre for the Living Planet at Imperial College London. We thank the Imperial College research computing services (High Performance Computing) for the usage of their resources. DPT acknowledges support from the Jarislowsky Foundation and NSERC. MH acknowledges support from the KR Foundation.

## Supplementary material

Supplementary material is made available compiled into a separate PDF file. The only empirical data used in this study was collated from third-party sources that are already publicly available.

