## Supplementary Information for "Species Richness and Speciation Rates for all Terrestrial Animals Emerge from a Synthesis of Ecological Theories"

#### Contents

**1 Analytical solution of species richness for protracted speciation** **1**

**2 Approximating spatial simulation richness with non-spatial analytical solution** **2**

**3 Numerical solutions for speciation-initiation rates** **3**

**4 Speciation rates: comparison between diet classes** **4**

**5 Speciation and Generation time power-laws** **5**

**6 Ectotherm abundances** **6**

#### 1 Analytical solution of species richness for protracted speciation

In the standard neutral model with point-mutation (PM) speciation (Etienne and Alonso, 2005), the expected number of species with abundance $n$ in a random sample of $J$ individuals from a metacommunity with $J_M$ individuals is given by:

$$E[S_n^{(PM)}|\theta, J] = \frac{\theta}{n}\frac{\Gamma(J+1)}{\Gamma(J+1-n)}\frac{\Gamma(J+\theta-n)}{\Gamma(J+\theta)} \tag{S1}$$

where $\theta = \dfrac{\nu(J_M-1)}{(1-\nu)}$ is the biodiversity number and $\nu$ is the speciation rate. The same expression would appear for the relative species abundance (RSA) in a spatially-implicit neutral model without dispersal limitation ($m = 1$). From Eqn. S1 it follows that the total expected number of species for this sample $E[S^{(PM)}|\theta, J] = \sum_{n=1}^{J} E[S_n^{(PM)}|\theta, J]$ can be put under the form (Chisholm et al., 2016):

$$E[S^{(PM)}|\theta, J] = \theta\left[\psi_0(\theta+J) - \psi_0(\theta)\right] \tag{S2}$$

where $\psi_0$ is the digamma function.

If instead we consider protracted (PROT) speciation in this neutral model, the expected number of species with abundance $n$ for a sample of $J$ individuals is given by (Rosindell et al., 2010):

$$E[S_n^{(PROT)}|\theta, J] = \frac{J!}{(J-n)!}\frac{\theta}{n}\left(\frac{\Gamma(J+\frac{\theta\beta}{\theta+\beta}-n)}{\Gamma(J+\frac{\theta\beta}{\theta+\beta})} - \frac{\Gamma(J+\beta-n)}{\Gamma(J+\beta)}\right) \tag{S3}$$

where $\beta = \dfrac{J_M - 1}{1+\tau}$ and now $\theta = \dfrac{\mu(J_M-1)}{(1-\mu)}$, with $\mu$ the speciation-initiation rate.

Note that if we use $\dfrac{J!}{(J-n)!} = \dfrac{\Gamma(J+1)}{\Gamma(J+1-n)}$, then Eqn. S4 can be rewritten as:

$$E[S_n^{(PROT)}|\theta, J] = \theta\left(\frac{1}{n}\frac{\Gamma(J+1)}{\Gamma(J+1-n)}\frac{\Gamma(J+\frac{\theta\beta}{\theta+\beta}-n)}{\Gamma(J+\frac{\theta\beta}{\theta+\beta})} - \frac{1}{n}\frac{\Gamma(J+1)}{\Gamma(J+1-n)}\frac{\Gamma(J+\beta-n)}{\Gamma(J+\beta)}\right), \tag{S4}$$

which corresponds to the subtraction of two terms identical to Eqn. S1, substituting the parameter $\theta$ for $\dfrac{\theta\beta}{\theta+\beta}$ and $\beta$, respectively.

Thus, as Eqn. S5 follows from Eqn. S1, then it follows that the total expected number of species for a sample of $J$ individuals should be given by:

$$E[S^{(PROT)}|\theta, J] = \theta\left\{\left[\psi_0\left(\frac{\theta\beta}{\theta+\beta}+J\right) - \psi_0\left(\frac{\theta\beta}{\theta+\beta}\right)\right] - [\psi_0(\beta+J) - \psi_0(\beta)]\right\}. \tag{S5}$$

### 2 Approximating spatial simulation richness with non-spatial analytical solution

As described on the Methods section, the analytical solution for the non-spatial neutral model (equation 0.1) is used in conjunction with the empirical richnesses and the global abundances obtained from the Madingley model to estimate a value of speciation-initiation for a given guild.

The main assumption to use this analytical solution is that, if one is concerned with the global richness for a given group, then dispersal or spatial heterogeneities in abundances or demographic rates can be disregarded in a first approximation.

To test this assumption we have run neutral simulations with protracted speciation using, for each body-size guild of terrestrial endotherms, data probed from the Madingley model for abundances, dispersal rates, and turn-over rates. Using the package pycoalescence (Thompson et al., 2020) to simulate spatial neutral simulations with protracted speciation (Rosindell et al., 2010) for different values of the time for speciation completion $\tau$. We then checked how the simulated global richness compared to the analytical richness, using the equation for the non-spatial model.

For some of the terrestrial endotherms' guilds, the global abundances probed from Madingley reach order $10^{10}$ individuals, making it computationally unfeasible to run a neutral simulation with all the individuals in the world. To solve this, we take random samples of these individuals as inputs for our neutral simulations. For each simulation, a fraction of $f = 10^{-4}$ of the global abundance for that guild is considered.

Thus, to compare the simulated richness with the analytical richness, we need an equation that gives the richness for a random sample of the individuals in a given community. The expected species richness $S$ for a random sample of $J$ individuals in a metacommunity with $J_M$ individuals will be:

$$S = \theta \left\{ \left[ \psi_0 \left( \frac{\theta\beta}{\theta+\beta} + J \right) - \psi_0 \left( \frac{\theta\beta}{\theta+\beta} \right) \right] - \left[ \psi_0(\beta + J) - \psi_0(\beta) \right] \right\}, \tag{S6}$$

according to equation S5. As before, $\beta = \frac{J_M - 1}{1+\tau}$, and $\theta = \frac{(J_M - 1)\mu}{(1-\mu)}$. Note that when this expression is used to estimate the speciation-initiation rates for endotherms in the main text (equation 0.1), we assume $J = J_M$ (or $f = 1$), since we use the global empirical richness, thus accounting for the totality of individuals.

The results for the comparison between simulated and analytical richnesses, for three different values of speciation-initiation rate $\mu$ are shown in Fig. S1:

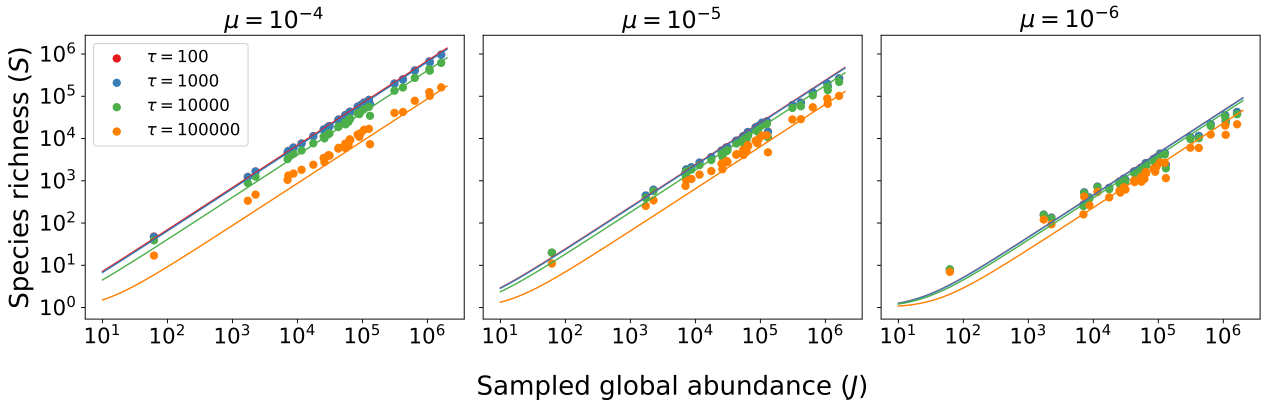


Figure S1: Comparison between simulated richness (circles) and analytical richness (Eqn. S5), for four different values of $\tau$, as a function of the number of individuals in a given sample. Each circle correspond to the results of one spatially explicit neutral simulation using abundances and demographic rates probed from the Madingley model for a given guild. For each simulation, we used a fraction sample of $f = 10^{-4}$ of the global abundance for each guild of terrestrial endotherm. For each plot, a different value of the speciation-initiation rate $\mu$ was used.

### 3 Numerical solutions for speciation-initiation rates

In Fig. 3 of the main text, the absence of speciation-initiation estimates for some of the guilds (those with smallest body sizes and largest values of $\tau$) is related to the lack of numerical solutions for $\mu$ at equation 0.1, for the corresponding empirical richness and Madingley abundance for that group. However, in some cases, not all of the 15 Madingley sampled abundances result in an impossibility to find a numerical solution for $\mu$.

If equation 0.1 is plotted as a function of $\mu$ alongside the value for the empirical richness for that group, the solution will correspond to the value of $\mu$ where the two curves intersect. In Fig. S2 we plot these curves for two values of abundance, corresponding to different Madingley samples of terrestrial herbivores, with body mass equal to 36 g, for $\tau = 100000$. For this group the global empirical richness is equal to 301.

As shown in Fig. S2(a), no biologically feasible speciation-initiation rate is possible for a group of $2.6 \cdot 10^7$ individuals to present 301 species for equilibrium richness, considering a protracted neutral model with $\tau = 100000$ generations for speciation completion. In contrast, if the number of individuals is $1.7 \cdot 10^8$ then such biologically feasible speciation-initiation rate can be obtained, as shown in Fig. S2(b). The value $\mu \approx 10^{-6}$ is the speciation-initiation rate where the empirical (red curve) and analytical (blue curve) richnesses intersect.

When we exclude only the Madingley samples for which no value of speciation-initiation rate can be obtained and average over those that can, additional estimates for these rates are

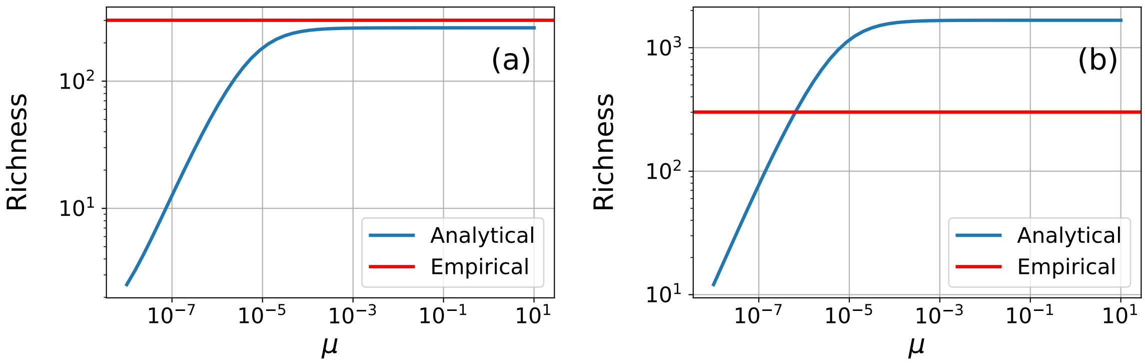


Figure S2: Plots of empirical richness and analytical richness, for two sampled abundances from Madingley with (a) $2.6 \cdot 10^7$ and (b) $1.7 \cdot 10^8$ individuals. The samples correspond to a guild of terrestrial herbivores with body mass equal to 36 g and corresponding global empirical richness equal to 301. For both plots, we use $\tau = 100000$ for the speciation-completion time.

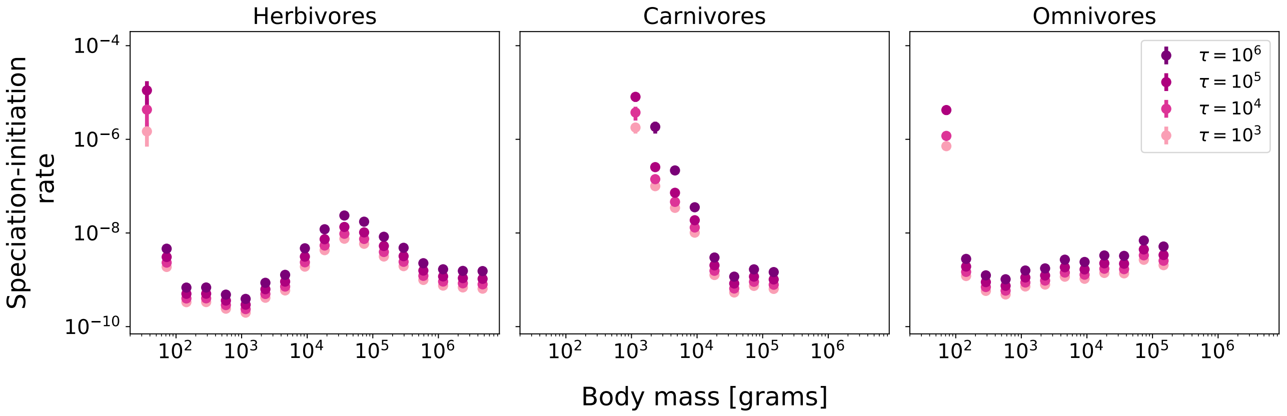


Figure S3: Numerical estimates for the speciation-initiation rates corresponding to Fig. 3 of the main text. The additional circles in this figure are averages of speciation-initiation rates excluding samples for which a solution for $\mu$ could not be found.

shown, in comparison to Fig. 3 of the main text. The additional estimates can be seen in Fig. S3. The new estimates appear at the first (for herbivores and omnivores) and up to the second (for carnivores) body size intervals, for values of $\tau$ greater than or equal to $10^5$. For the three diet classes, the values corresponding to the first body size interval and $\tau = 10^6$ are not shown, since no solutions for $\mu$, for any of the 15 samples, could be found.

### 4 Speciation rates: comparison between diet classes

The values obtained for speciation rates for endotherms (per species per million yesars) as functions of body mass, shown in Fig. 5 in the main text, can also be plotted separately for each value of speciation-completion time, $\tau$ (Fig. S4). These plots allow for a clearer comparison of speciation rates between the three diet classes. Patterns of decrease for speciation rates as functions of body mass for herbivores, carnivores and omnivores are qualitatively similar for each value of $\tau$, showing the same trend of decrease for increasing $\tau$. As discussed in the main text, while speciation rates are similar herbivores and omnivores for the same body size group, carnivores present higher rates for values of body mass lower than $10^4$. For body masses higher than $10^4$, carnivores and herbivores present similar values of speciation rates, for all values of $\tau$, while omnivores show slightly smaller values.

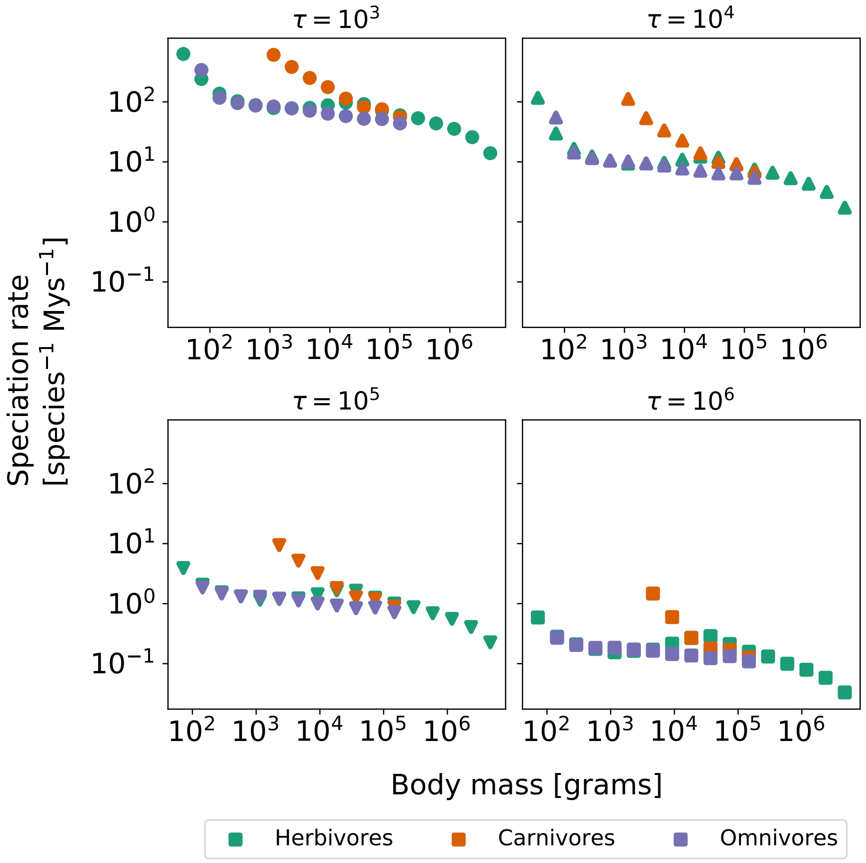


Figure S4: Comparison of endotherms' speciation rates (per species per million years) between herbivores, carnivores and omnivores as functions of body mass, for each value of $\tau$.

### 5 Speciation and Generation time power-laws

Here we show the fitted functions for generation times S5 and speciation rates S6 using in the main text as part of the pipeline of estimation of global richnesses. Best fit for power-law function $f(x) = Ax^b$ is shown for each case with 95% confidence bands. For generation times, a single model was fitted grouping data for herbivores, carnivores and omnivores, while for speciation rates each diet class and each value of $\tau$ was fitted separately.

Parameters for the best fit for generation times are $A = 8.9688 \cdot 10^{-8} \pm 7.1442 \cdot 10^{-9}$ and $b = 0.17089351 \pm 0.00650863$. For speciation rates, fitted parameters are show in table S6.

Table S6: Parameters for fitted power-law functions of speciation rates per species per million years for endotherms (Fig. S6).

| $\tau$ | Diet class | A | b |
|---|---|---|---|
| $10^5$ | Herbivore | $5.10146851 \pm 1.54581723$ | $-0.17243167 \pm 0.02404888$ |
| | Carnivore | $91.9436859 \pm 80.8498805$ | $-0.39902811 \pm 0.08314994$ |
| | Omnivore | $2.91456643 \pm 0.18320385$ | $-0.11594684 \pm 0.00676674$ |
| $10^6$ | Herbivore | $0.61139078 \pm 0.18455592$ | $-0.16084877 \pm 0.02402343$ |
| | Carnivore | $6.65143951 \pm 6.65313716$ | $-0.33687969 \pm 0.09369326$ |
| | Omnivore | $0.37119641 \pm 0.03045594$ | $-0.10146528 \pm 0.00898029$ |

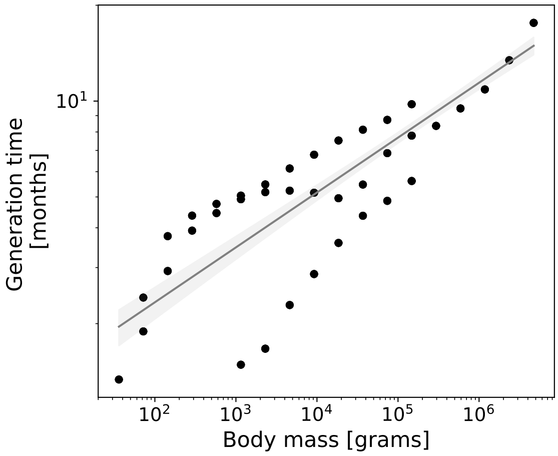


Figure S5: Power-law fitted function ($Ax^b$) for generation times for Madingley endotherms. Data for herbivores, carnivores and omnivores were grouped to fit a single model. Values of the parameters for the best fit are $A = 8.9688 \cdot 10^{-8} \pm 7.1442 \cdot 10^{-9}$ and $b = 0.17089351 \pm 0.00650863$.

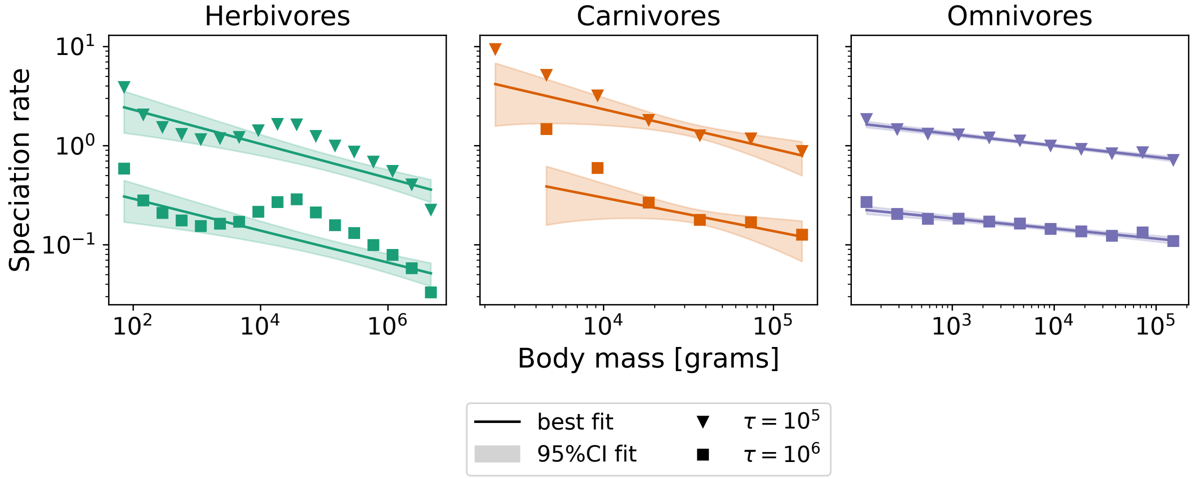


Figure S6: Power-law fitted functions ($Ax^b$) for speciation rates per species per million years relationship with body size for endotherms. Separate models were fitted for each diet class and each value of $\tau$. Parameters for the best fit in each case can be seen in table S6.

### 6 Ectotherm abundances

Similarly to the patterns for endotherms (Fig. 2 in the main text), herbivores and omnivores show a decreasing trend with body mass, both for semelparous (first row - Fig. S7) and iteroparous ectotherms (second row - Fig. S7). For carnivores, abundances peak at 10 $g$ and $10^3$ $g$ for semelparous and iteroparous ectotherms, respectively. Semelparous carnivores present a slight tendency of decreasing abundances at larger body masses, while a more pronounced tendency can be seen for iteroparous carnivores at larger body masses.

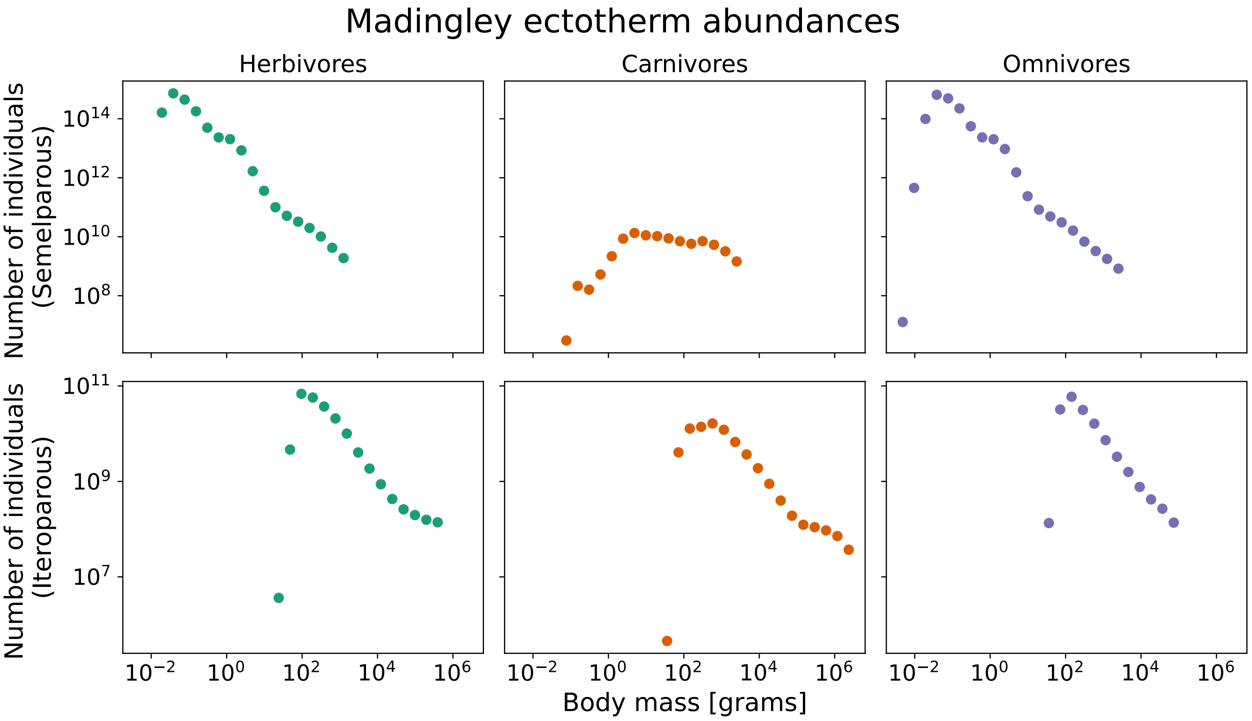


Figure S7: Average global abundances (in number of individuals) from the Madingley model at equilibrium for terrestrial ectotherms as functions of body mass for three diet classes: herbivores, carnivores and omnivores. First row corresponds to semelparous ectotherms and second row corresponds to iteroparous ectotherms.


Ecology letters 8:1147–1156.

Rosindell, J., S. J. Cornell, S. P. Hubbell, and R. S. Etienne. 2010. Protracted speciation revitalizes the neutral theory of biodiversity. Ecology Letters 13:716–727.

Thompson, S. E., R. A. Chisholm, and J. Rosindell. 2020. pycoalescence and rcoalescence: Packages for simulating spatially explicit neutral models of biodiversity. Methods in Ecology and Evolution 11:1237–1246.
